# Increased speech representation in older adults originates from early response in higher order auditory cortex

**DOI:** 10.1101/294017

**Authors:** Christian Brodbeck, Alessandro Presacco, Samira Anderson, Jonathan Z. Simon

**Affiliations:** Institute for Systems Research, University of Maryland, College Park, Maryland; Department of Otolaryngology, University of California, Irvine, California; Department of Hearing and Speech Sciences, University of Maryland, College Park, Maryland; Department of Electrical and Computer Engineering, University of Maryland, College Park, Maryland; Department of Biology, University of Maryland, College Park, Maryland

## Abstract

Previous research has found that, paradoxically, while older adults have more difficulty comprehending speech in challenging circumstances than younger adults, their brain responses track the acoustic signal more robustly. Here we investigate this puzzle by using magnetoencephalography (MEG) source localization to determine the anatomical origin of this difference. Our results indicate that this robust tracking in older adults does not arise merely from having the same responses as younger adults but with larger amplitudes; instead they recruit additional regions, inferior to core auditory cortex, as part of an early response peak at ~ 30 ms relative to the acoustic signal.

## 2 Introduction

While older adults have more difficulty comprehending speech in challenging circumstances, their brain responses track acoustic speech signals more robustly than the brain responses of young adults [1,2]. There have been several candidate explanations put forward to account for this observation.

Older adults exhibit increased response amplitudes to even simple tone stimuli [3,4], suggesting that enhanced speech tracking could be due to amplified representations of any auditory features. This might arise from central compensatory gain mechanisms that restore the representation of sounds at the cortical level despite degraded auditory brainstem responses [5,6]. Animal models also suggest that aging may alter the balance between excitatory and inhibitory processes (decreasing inhibition) in the cortex, acting at several levels along the auditory pathway [7–12], leading to stronger cortical stimulus-driven responses.

On the other hand, there is also evidence for a reorganization of task-dependent networks, in which older adults recruit additional higher order cortical regions to compensate for age-related changes in lower order processing [13,14]. Aging may also compromise the efficient use of cognitive resources because of decreased cortical network connectivity [13], leading to redundant processing across areas.

Consequently, increased neural speech tracking might arise from several different underlying changes. These changes can be distinguished in whether they uniformly affect responses that are also involved in young adults, or whether they involve increased recruitment of additional regions. Here we investigate increased speech representation by using source localization to determine the anatomical source of the difference. Our results indicate that robust tracking in older adults does not arise merely from having the same responses as younger adults with larger amplitudes, but instead from recruiting additional regions, ventral to core auditory cortex, for processing speech.

## 3 Methods

MEG data were collected from a sample of 17 younger (18-27 yr, 3 men) and 15 older adults (61-73 yr, 5 men) with clinically normal hearing, described in detail in [1]. Here we only analyze data from participants listening to two one-minute long segments of an audiobook rendition of *The Legend of Sleepy Hollow* (https://librivox.org/the-legend-of-sleepy-hollow-by-washington-irving). Each segment was repeated 3 times for a total of 6 minutes of data per subject.

For details on the basic MEG data analysis see [15]; here we focus on deviations from the procedure described there. Raw data were filtered between 1-8 Hz [1], downsampled to 100 Hz and projected to virtual current dipoles distributed on the cortical surface. These localized brain responses were individually modeled as driven by the analytic envelope of the acoustic stimulus, using a linear filter model [16]. Boosting was used to estimate the optimal filter known as the Temporal Response Function (TRF) [17]. Response functions were generated from a basis of 50 ms Hamming windows, distributed at 10 ms intervals in the kernel window of 0 – 500 ms. Thus, each response function was described as the sum of 50 scaled Hamming windows with scaling values determined by boosting.

Model fits were evaluated based on the *z*-transformed Pearson correlation between predicted and actual source localized responses. To determine the predictive power of the model with bias correction, we subtracted from the correct model fit the model fit obtained from using a misaligned version of the same stimulus (obtained by switching the first and second half of the acoustic stimulus). Since the goal of this study was to analyze brain responses associated with time-locked auditory processing, the analysis was restricted to the temporal lobes of both hemispheres.

To localize the source of the higher predictability of older adults’ brain responses, model fits were compared between the two groups by performing *t*-tests at each virtual current dipole, while controlling for multiple comparisons using threshold-free cluster enhancement (TFCE) [18]. To statistically assess hemispheric differences, results from the right hemisphere were mapped to the left hemisphere as described in [15], and left-right difference maps were then compared between groups.

The area of significant group differences was then taken as the region of interest (ROI) in which to analyze estimated response functions, and, specifically, to determine the latency of the responses contributing to the effect. For subsequent analyses, absolute values of the response functions were used to avoid an influence of arbitrary differences in current direction due to cortical surface orientation. The mean absolute value over source dipoles within the ROI was extracted for each time point of the response function. The result was an estimate of the magnitude of the response’s contribution to predictions at each time point. This time course exhibited characteristic peaks that were used to determine the latency at which older adults’ responses in the ROI were stronger than young adults’ responses. A one-tailed *t*-test was performed at each time point, again corrected using TFCE.

Finally, to confirm that the difference in responses in the peak thus identified was indeed specific to the anatomical region identified for enhanced speech representation, the average absolute response during the peak was extracted for each dipole. The spatial distribution of this peak response was then compared between groups, again corrected with TFCE.

## 4 Results

Brain responses of older adults were predicted significantly better than responses of young adults in a region of the left temporal lobe clearly outside core auditory cortex (Figure 1A). Though this effect was not significant in the right hemisphere, there was no significant difference between hemispheres either (*p* = .393). Group averaged model predictions suggest that for both groups, predictions were best near core auditory cortex (Figure 1B). In older adults, however, the area in which the model made good predictions extended further. The fact that the spatial peak of the difference is located inferior to the spatial peak of the group average indicates that this is a true difference in neural source locations, rather than an artifact due to an amplitude difference combined with spatial dispersion of MEG source estimates, which would instead manifest as a difference map peak close to the average peak.

**Figure 1:**
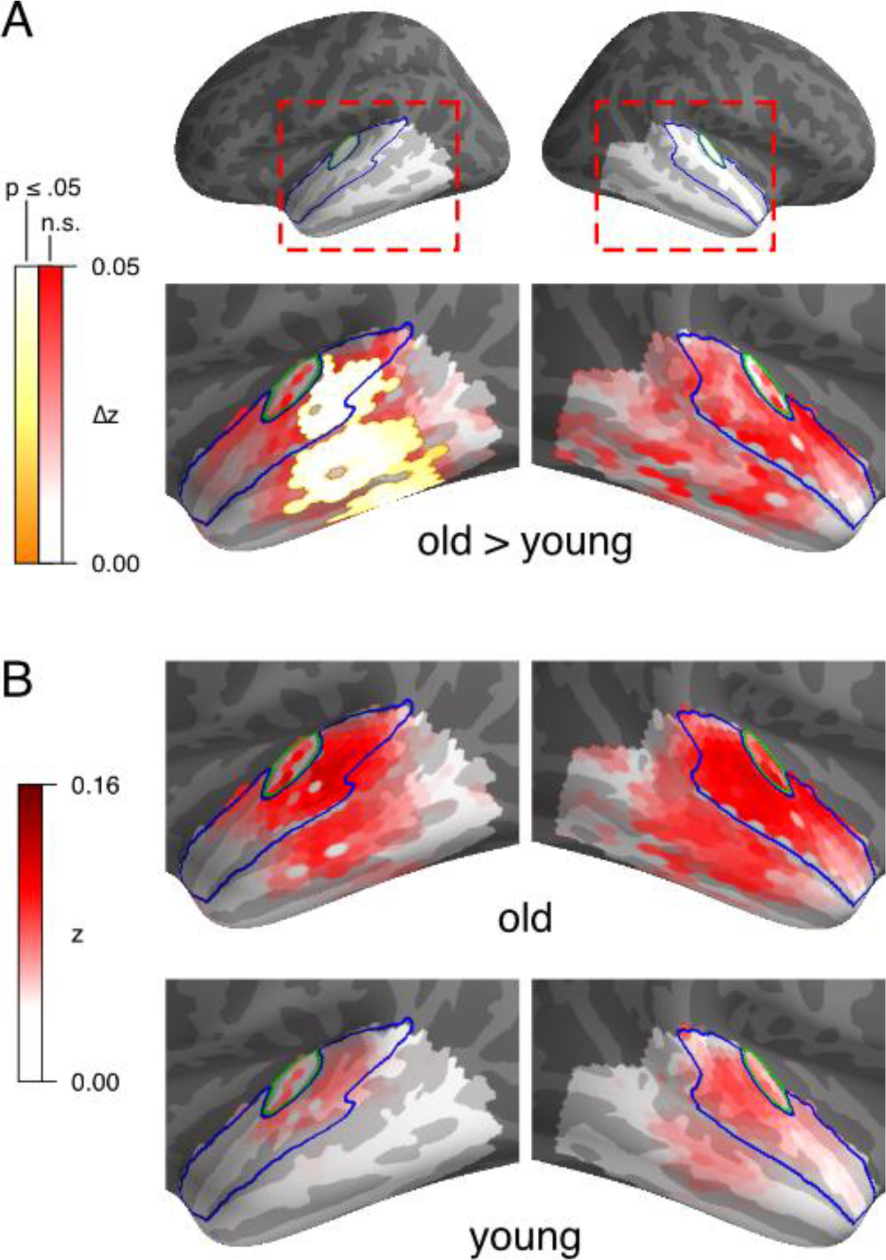
Model fit, expressed as z-transformed Pearson correlation between predicted and measured (source transformed) responses. A) Significantly better predicted brain responses in older adults than younger adults, in a region below left core auditory cortex (p ≤ .05 corrected within indicated bilateral temporal lobe). Outlines indicate Heschl’s gyrus (core auditory cortex, green) and superior temporal gyrus (blue). B) Average model prediction quality for each participant group separately.

The response function within the significant area exhibited characteristic peaks around 30 and 100 ms [15,19], with an additional peak around 180 ms for older adults (Figure 2A). A significant difference between older and younger adults emerged in the earliest response peak (TFCE time course significant 10-20 ms, *p* = .029, one-tailed). No other peak was significantly different, even when excluding the data encompassing the first peak up to 70 ms to implement a step-down procedure [20].

**Figure 2:**
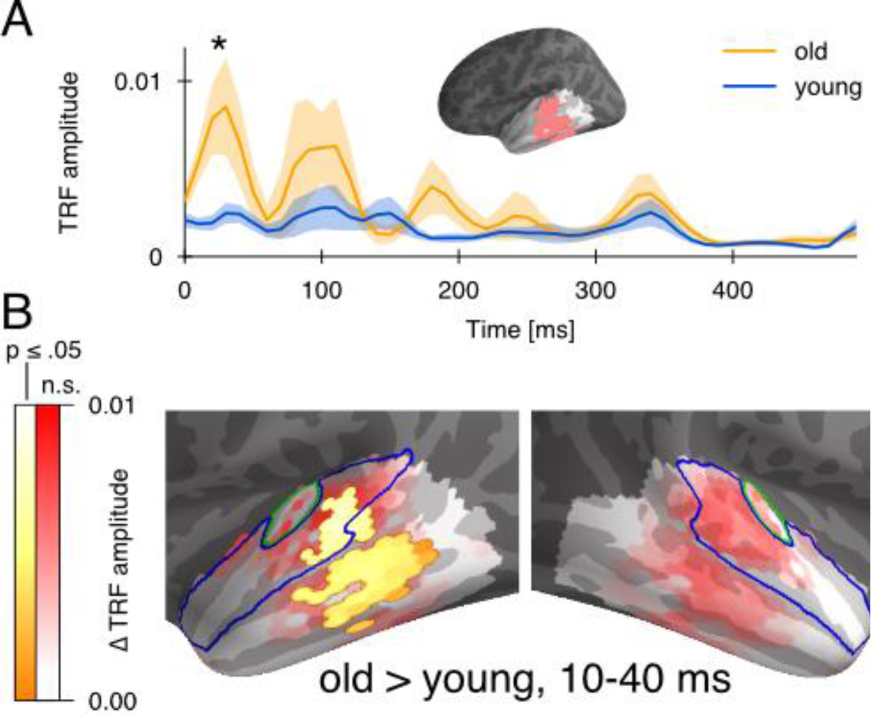
Response function in the region of difference. A) Mean absolute response function in the ROI based on significant difference in model fit (cf. Figure 1), with standard error. Older adults had significantly higher amplitude in the earliest response peak. B) Anatomical distribution of the difference in the absolute response strength during the window defined based on the first peak (10-40 ms). The anatomical distribution closely resembles the region of the difference in model fit.

The spatial distribution of the amplitude difference in the first peak, shown in Figure 2B, closely resembled the spatial extent of the model fit difference (Figure 1A). This confirms that the difference in response magnitude in the first peak was indeed specific to the region with improved model fit. Thus, enhanced representation of the acoustic speech signal in older adults has significant contributions from an increased early response in higher order auditory cortex.

## 5 Discussion

Compared to young adults, older adults’ brain responses more accurately track amplitude modulations in the acoustic envelope of speech in the 1-8 Hz range. Source localized MEG responses suggest that this effect is particularly pronounced in the left temporal lobe, in a region lateral and inferior to primary auditory cortex. This suggests that older adults, rather than exhibiting the same responses as young adults with uniformly higher response amplitudes, disproportionately recruit higher order auditory cortex during speech perception.

Analysis of the response functions suggested that the largest contribution to this effect occurred in the earliest response peak with only about 30 ms latency. This peak is associated with processing of acoustic stimulus properties, as opposed to a later peak around 100 ms which is more sensitive to attended than unattended acoustic features [19]. This suggests that in older adults, the initial stage of cortical speech sound processing engaged a larger neural population in higher order auditory regions.

This is consistent with several studies reporting that aging might alter the balance between inhibitory and excitatory neural mechanisms in the cortex [7,10–12]. The resulting increase in neural excitability and decrease in neural selectivity would lead to larger onset responses to an auditory signal [21]. On the other hand, this early overrepresentation might also be an indication of a disproportional allocation of attentional resources to a task that is perceived as more difficult [22], as suggested by several studies showing that older adults perform more poorly than younger adults during a dual task (e.g. auditory and memory) [23–25].

Figure 2A suggests that older adults also exhibit a larger peak than younger adults around 180 ms. While this difference was not significant in this analysis, the difference does reach significance when the ROI is enlarged to include the temporal lobes of both hemispheres. This enhanced later amplitude is consistent with improved stimulus reconstruction seen for older but not young adults when time windows longer than 150 ms are used [2].

A recent large scale investigation suggests that age-related changes in simple tone-evoked magnetic fields are characterized by a cumulative delay, which is associated with decreased grey matter volume in higher order auditory cortex [3] (the relation to increased amplitudes was not analyzed there). This region (delimited with the greater spatial confidence of MRI) lies within the region found here to track the acoustic stimulus with greater fidelity in early responses of older adults.

Together, these results suggest that altered auditory processing in older adults might be related to changes in early responses in higher order auditory areas. These may arise from a variety of phenomena, including degraded network communication, imbalance between inhibition and excitation, and inefficient use of cognitive resources.

